# A mathematical model of the Alzheimer’s Disease biomarker cascade demonstrates statistical pitfalls in identifying neurobiological surrogates of cognitive reserve

**DOI:** 10.1101/2023.10.26.563793

**Authors:** Florian U. Fischer, Susanne Gerber, Oliver Tüscher, the Alzheimer’s Disease Neuroimaging Initiative

## Abstract

**Introduction:** In order to investigate neurobiological surrogates of cognitive reserve, statistical interaction analyses have been put forward and used by several studies. However, as these neurobiological surrogates are potentially affected by neurodegeneration as part of the amyloid cascade, which is characterized by chronological time-moderated associations between biomarkers, cross sectional sampling in combination with the disregard of time as a confounder could introduce interaction effects that may be misinterpretabed as cognitive reserve in statistical analyses.

**Methods:** We modeled the amyloid cascade with a minimal set of three biomarkers amyloid load, corticospinal fluid tau, hippocampal volume and cognitive outcome using a differential equation system, whose parameters were estimated from empirical data from the ADNI. Interaction effects between pathology markers amyloid and tau with hippocampal volume as potential marker of cognitive reserve were estimated on two simulated data samples. Both samples were calculated from varying amyloid, tau and hippocampal volume for the initial configuration of individual trajectories. For Sample 1, data points were sampled at a fixed time after baseline. For Sample 2, data points were sampled at random time points.

**Results:** Regression analyses on Sample 1 yielded estimates for interaction effects of 0. For Sample 2, estimates were -.1692 and -.0807 for amyloid and tau with hippocampal volume, respectively. The interaction effect estimates for Sample 2 decreased several orders of magnitude when taking into account the timepoint of sampling.

**Conclusion:** Studies aiming to investigate neurobiological surrogates of cognitive reserve that are affected by Alzheimer’s Disease-related neurodegenerative processes need to consider inter-individually varying sampling time points in the data to avoid misinterpreting interaction effects.

## Introduction

Individual trajectories of cognitive decline demonstrate considerable variance in the presence of comparable amounts of neuropathology associated with age-related neurodegenerative diseases, most commonly Alzheimer’s Disease (AD). This has caused the exertion of considerable effort to conceptualize resilience to cognitive decline and its potential mechanisms as well as the identification and quantification of neurobiological bases thereof [1, 6, 14, 17, 28, 29, 34]. According to the predominant concepts and nomenclature, the phenomenon of resilience has been trichotomized into brain maintenance, brain reserve and cognitive reserve [29]. Briefly, brain maintenance refers to the individually varied accumulation of age-related brain pathology. Brain reserve represents a quantifiable neurophysiological state of brain components, whose depletion over time by neuropathological processes to levels inducing cognitive impairment takes longer in individuals with higher starting quantities. Cognitive reserve on the other hand represents some form of functional adaption to the presence of neuropathology, modulating its impact on cognition. As the direct observation of cognitive reserve may be exceedingly difficult from an experimental design point of view, surrogates that refer to the neurophysiological prerequisites of cognitive reserve, such as e.g. white matter or regional grey matter volume etc., may be investigated. To this end, cognitive reserve has been operationalized statistically as a moderation effect of measures of pathology on cognition by reserve factors [29].

However, as AD acts on many structures of the brain, such neurophysiological surrogates of cognitive reserve have a high chance of being affected as well, and thus arguably represent a convolute of effects of the disease process on the brain and information referring to the physiological basis of cognitive reserve. This raises the question of how the concepts of resilience are represented within the consensus model of AD.

Extensive research has produced a biomarker cascade model for AD, wherein biomarkers are altered consecutively in a fixed order [16]. The model includes cognitive outcome as its endpoint as well as biomarkers for molecular pathology, i.e. amyloid and tau, and biomarkers referring to neural injury to brain components such as hippocampal volume. Although findings of imaging studies from the last decade have considerably expanded our knowledge of the temporo-spatial associations of imaging biomarkers in the course of AD [31], this basic structure of the biomarker cascade model remains at the core of current conceptions of AD [15].

This model incorporates cognitive reserve in the sense that the rate of change of cognitive outcome over time varies relative to biomarkers based on cognitive reserve. However, the possibility that some biomarkers may refer not only to neuronal injury due to AD but also to brain reserve or cognitive reserve, as, e.g., hippocampal volume [34], is not explicitly incorporated in the biomarker cascade model. If one argues that the cascade character of the model derives from time-lagged associations between the consecutively altered biomarkers where the rate of change of a later altered biomarker depends on the level of an earlier altered biomarker, accommodating initially varying individual biomarker configurations (representing brain maintenance or brain reserve) would simply require a corresponding shift of the biomarkers along the time axis. In addition, if the biomarkers contribute independently to the rate of cognitive decline, that rate would have to depend on the biomarkers accordingly.

However, under the premise of this model, the identification of surrogates of cognitive reserve via statistical interaction effects [29] may be problematic. Consider, for the sake of argument, a group of model subjects whose biomarker state trajectories are described by the model from the previous paragraph, where the initial configuration of biomarkers for the subjects varies randomly and independently due to brain maintenance or brain reserve. We can make the following observations for the data points along these trajectories: (1) cognitive outcome will be lower at later time points, as the level of cognitive outcome depends on the amount of time it has been subject to the rate of change that is a function of biomarker levels. (2) all biomarkers themselves will also generally and thus jointly be more altered at later time points despite initially random distribution. If one were to randomly sample one data point from each trajectory to build a data sample, as is done in cross-sectional studies, and conduct statistical interaction analyses between a pair of biomarkers, one would find the following due to observations (1) and (2): data points, where only one of the main effect variables is altered will demonstrate less altered cognitive outcome, as they correspond to earlier time points from the trajectories. Data points, where both main effect variables are altered, will demonstrate more altered cognitive outcome as they correspond to later time points from the trajectories. A statistical analysis on such a sample is likely to report an interaction effect between biomarkers, which is an artifact produced by the underlying temporal dynamics of the biomarker trajectory, their initial variation and the sampling procedure but could be mistakenly interpreted as an indicator of cognitive reserve [29].

In order to test this hypothesis, we estimated a minimal mathematical model from empirical longitudinal data that formalizes the biomarker cascade model, accommodates individually differing initial biomarker configurations and strictly additive, i.e. independent, effects of biomarkers on the rate of change of cognitive outcome. We then estimated regression models on the simulated data produced by the model, investigating interaction effects between biomarkers of molecular pathology, cerebral amyloid beta and corticospinal fluid tau, and hippocampal volume, a biomarker that may partially refer to cognitive reserve surrogates [34]. The findings of this study may inform the experimental design and interpretation of studies investigating neurophysiological surrogates of brain reserve and cognitive reserve.

## Materials and Methods

### Data description

Data used in the preparation of this article were obtained from the Alzheimer’s Disease Neuroimaging Initiative (ADNI) database (`adni.loni.usc.edu`). The ADNI was launched in 2003 as a public-private partnership, led by Principal Investigator Michael W. Weiner, MD. The primary goal of ADNI has been to test whether serial magnetic resonance imaging (MRI), positron emission tomography (PET), other biological markers, and clinical and neuropsychological assessment can be combined to measure the progression of mild cognitive impairment (MCI) and early Alzheimer’s disease (AD). For up-to-date information, see www.adni-info.org.

### Subjects

For parameter estimation of the mathematical model, we included subjects of the ADNI cohort that had at least two data points with available Florbetapir (AV45) amyloid-PET summary scores, biospecimen results for corticospinal fluid (CSF) tau, available hippocampal volumetry as well as Alzheimer’s Disease Assessment Scale cognitive outcome scores (ADAS-cog). Additionally, subjects were only included if their amyloid-PET summary score was greater or equal to 1.1 at least once. This was to ensure that data points would represent the continuum of the AD-type neurodegenerative trajectories. Demographical data are summarized in Table 1.

**Table 1.** Demographical data. This table summarizes the demographical data of the sample used to estimate the mathematical model parameters. Values represent status at the earliest available time point. Age and education in years, data time in months. MCI, mild cognitive impairment. Mean *±* SD.

| Status | N | female | Age | Education | Data time span |
| --- | --- | --- | --- | --- | --- |
| Preclinical AD | 52 | 32 | 74.3±5.5 | 16.4±2.6 | 90.1±38.1 |
| MCI due to AD | 111 | 50 | 71.6±6.4 | 16.1±2.8 | 72.2±33.0 |
| AD | 13 | 5 | 76.7±7.0 | 15.4±3.0 | 39.7±28.0 |
| Total sample | 176 | 87 | 72.8±6.4 | 16.1±2.7 | 75.1±36.4 |

### Neuropsychological assessment

In general, subjects underwent neuropsychological assessment every 12 months over a span of years varying individually. Details regarding the cognitive assessment within the ADNI are given in the publicly available procedures manual: https://adni.loni.usc.edu/wp-content/uploads/2008/07/adni2-procedures-manual.pdf. Within the scope of this study, the cognitive Alzheimer’s Disease Assessment Scale (ADAS-cog), which spans several cognitive domains [25] was included. The ADAS-cog scale is oriented such that higher scores indicate worse cognitive performance.

### Corticospinal fluid measurement

CSF biospecimen were stored and analyzed at the Penn ADNI Biomarker Core Laboratory at the University of Pennsylvania, Philadelphia. The multiplex xMAP Luminex platform (Lumnix Corp., Austin, TX) was used to measure CSF concentrations of total tau. For the present study, total tau was included instead of phosphorylated tau, as the latter is included in the former. Total tau thus includes tauopathy specific to AD as well as more general age-related tauopathy that may be relevant for cognitive outcome [5]. A more detailed description of data collection and processing of the CSF samples can be found in [27] and at http://adni.loni.usc.edu/methods.

### Imaging data acquisition and processing

Inversion-recovery spoiled gradient recalled (IR-SPGR) T1-weighted imaging data were acquired on several General Electric 3T scanners using a gradient echo sequence with 11 degree flip angle and a voxel size of 1.0^2^ x 1.20mm^3^. AV45 amyloid PET imaging data were acquired on several types of scanners using different acquisition protocols. PET data underwent a standardized preprocessing procedure at the ADNI project to increase data uniformity. Imaging protocols and preprocessing procedures can be accessed at the ADNI website: http://adni.loni.usc.edu/methods/.

Using T1-weighted images, total intracranial volume (TIV) as well as hippocampal volume were calculated at ADNI core laboratories using published tissue segmentation methods [7, 8, 10]. Hippocampal volume was also normalized dividing by the total intracranial volume (TIV).

Subjects’ global cortical amyloid load was calculated from AV45 PET images according to ADNI procedures http://adni.loni.usc.edu/methods/pet-analysis/. To this end, cortical amyloid was calculated as the average of the AV45 uptake in the frontal, angular/posterior cingulate, lateral parietal and temporal cortices normalized by dividing by the mean uptake in the cerebellum [9].

### Development of a mathematical disease model

We model the consensus biomarker model of AD mathematically as a linear inhomogeneous differential equation model with constant coefficients. The individual biomarkers as well as the cognitive outcome are represented as system state vector 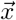, wherein the biomarkers are ordered inversely to their chronological order, i.e. each biomarker comes before the set of biomarkers that precede it in the cascade model. Specifically, the first entry is the cognitive outcome followed by hippocampal volume,

CSF tau and amyloid-PET. The associations among the biomarkers are represented by the triangular matrix *A*, and the association with time is represented by 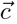. The matrix *A* is triangular due to the constraint on hypothetical causality implied by the chronological order of the biomarker cascade model, i.e. a biomarker can only conceivably depend on the biomarkers that precedes it in the cascade model. Hence also the ordering of the state vector 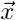. The model is thus represented by a differential equation system of the form.

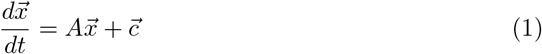

### Determining the model parameters

We determined the association of the *i* individual biomarkers with each other and with time using linear mixed effects models (see section statistical methods below for details). The estimated regression coefficients were inserted in equations representing the association of each biomarker *x*_*i*_ with time *t*, the biomarkers preceding it in the biomarker cascade *x*_*j*_ as well as the modulation of the association with time by the respective biomarkers *x*_*j*_*t*, where *j > i*. These equations take the form

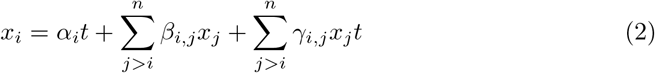

Their derivatives are

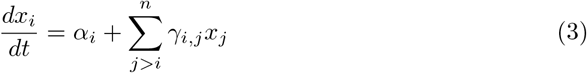

We can thus set the model parameters by inserting them into equation (1), setting the matrix *A*’s elements *a*_*i,j*_ = *γ*_*i,j*_ and the inhomogeneous component *c*_*i*_ = *α*_*i*_.

### Analytical solution

The solution to a linear inhomogeneous differential equation system has the form

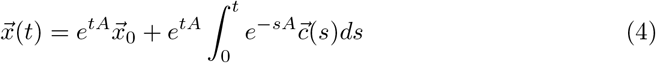

We can calculate the matrix exponential exactly using the Taylor series, which aborts after a finite number of terms as *A* is a triangular and thus nilpotent matrix.

Additionally, as 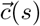 is time independent in our model, it can be simplified to 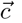. Substituting the matrix exponentials with their Taylor series expansion and calculating the integral then yields the solution to the model

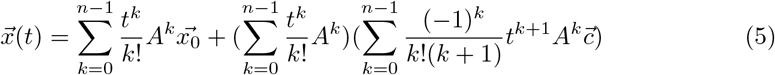

## Statistical methods

Empirical data was z-standardized throughout. In order to define the mathematical model parameters, we estimated for each biomarker and the cognitive outcome score as dependent variable a family of nested linear mixed effects models consisting of every possible combination of the biomarkers preceding the respective biomarker/the cognitive outcome score in the biomarker cascade as well as their interaction terms with time as predictor, as well as the model without other biomarkers. Time (in months after baseline), the covariates age and education (in years) as well as as a random intercept for each subject were included in all models. Using conditional Akaike Information Criterion [11, 30], the model best describing the data was then selected and its estimates used to set the model parameters as described above. If a preceding biomarker’s interaction term was not included in the best fitting model, its corresponding entry in *A* was set to zero.

In order to assess the correspondence of the simulated data produced by the mathematical model with the available empirical data points, we used the biomarker and cognitive outcome data of each subject at baseline (if available) to calculate the model predictions of cognitive outcome and biomarker statuses at subsequent time points that were also available in the real data. These simulated data points were then compared to the corresponding empirical data via simple linear regression models, with the mathematical disease model predictions as the predictor and the empirical data as dependent variable.

The aim of the main analysis was to demonstrate the emergence of interaction effects due to cross sectional sampling of data simulated under the premise of the AD biomarker cascade model (see above). To this end, we first simulated individual biomarker and cognitive status trajectories over time in the following way. Each biomarker was varied between -2 and +2 standard deviations (SD) from the mean of its empirical distribution with a step size of 0.1 SD in order to span the majority of the range of their empirical distribution. Each possible combination of the resulting biomarker statuses then yielded the set of all initial configurations, while cognitive outcome was set to its empirical mean in all of them. Subsequently, for each of these initial configurations, biomarker and cognitive outcome status were calculated for the time points that were measured in the empirical data - 6 to 186 months after baseline - using the mathematical model described above.

Second, we estimated regression models with cognitive outcome as dependent variable, the biomarkers as main effects and the interaction terms of amyloid-PET with hippocampal volume and CSF tau with hippocampal volume. The regression models were estimated on two subsets of the simulated individual trajectories described in the previous paragraph, one representing a cross-sectional analysis informed of the time-correspondence of data points, where a regression model was estimated for each calculated time point after baseline on the simulated data points corresponding to the respective time point (consistent sampling). The other represented a ‘naturalistic’ cross-sectional analysis uninformed of the time-correspondence, where one simulated data point was sampled for each individual trajectory at a random time point (random sampling). The random sampling and regression estimation was repeated 1000 times for this second analysis and estimated regression coefficients were averaged. This analysis was then repeated with the effect of time regressed out of each variable by using each variable’s residuals from a simple regression with the respective variable as dependent and time after baseline as independent variable (random sampling from residuals).

For each set of regression estimates, i.e. consistent sampling, random sampling and random sampling from residuals, we reported the 95% confidence interval and the average of regression coefficient estimates (see Table 3).

### Software packages

For the implementation of the mathematical model, simulation and statistical data analysis as described above, we used R version 4.1.2 with the following packages: psych version 2.2.9 [24], expm version 0.999-7 [18], stringr version 1.5.0 [33], cAIC4 version 1.0 [26], nlme version 3.1-155 [22, 23], remef version 1.0.7 [13], lme4 version 1.1-31 [2], Matrix version 1.5-3 [3], ggplot2 version 3.4.0 [32].

## Results

### Mathematical model parameters

For the model parameters *A* and 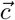 estimated by linear mixed-effects models, please refer to Table 2. Specifically, for the independent change of the biomarkers over time part of the model 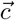, an increase for amyloid PET and CSF tau as well as a decrease for ADAS-cog and hippocampal volume over time were estimated from empirical data. For the associations among biomarkers, higher amyloid PET was associated with an increase of CSF tau and ADAS-cog as well as a decrease of hippocampal volume over time. Higher CSF tau was associated with increasing ADAS-cog over time. Finally, lower hippocampal volume was associated with increasing ADAS-cog over time.

**Table 2.** Estimated model parameters. This table summarizes the model parameters *a*_*i,j*_ ∈*A* and 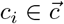 estimated by linear mixed effects models. The elements of *A* quantify how much the biomarkers in the rows of the table change depending on the biomarkers in the columns. The elements of 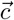 in the last column quantify how much the biomarkers in the rows of the table change depending on time. ADAS-cog, cognitive Alzheimer’s Disease Assessment Scale. HippoV, Hippocampal Volume. CSF tau, corticospinal fluid total tau. AV45, florbetapir amyloid PET score.

| $a_{i,j}$ | ADAS-cog | HippoV | CSF tau | AV45 | $c_i$ |
| --- | --- | --- | --- | --- | --- |
| ADAS-cog | 0 | -.23159 | .04549 | .11228 | -.08505 |
| HippoV | 0 | 0 | 0 | -.06495 | -.41949 |
| CSF tau | 0 | 0 | 0 | .01360 | .10513 |
| AV45 | 0 | 0 | 0 | 0 | .18910 |

**Table 3.** Regression estimates. This table summarizes standardized regression coefficient estimates, average and 95% confidence interval, on the simulated data from the mathematical model whose parameters are shown in Table 2. Consistent, data points were sampled cross sectionally from one corresponding timepoint after baseline from each individual trajectory (consistently across trajectories). Random, data points were sampled from a random timepoint cross sectionally from each individual trajectory. Random f. residuals, as in random sampling but with the variance associated with time after baseline removed from main effect variables. AV45, florbetapir amyloid-PET summary score. CSF tau, corticospinal fluid total tau. HippoV, hippocampal volume.

| Model term | Consistent | Random | Random f. residuals |
| --- | --- | --- | --- |
| AV45 | .3369 | .2627 | .2987 |
|  | [.3171,.3567] | [.2626,.2627] | [.2986,.2988] |
| CSF tau | .1679 | .1286 | .1448 |
|  | [.1665,.1692] | [.1286,.1287] | [.1447,.1449] |
| HippoV | -.8710 | -.7586 | -.7498 |
|  | [-.8731,-.8690] | [-.7587,-.7585] | [-.7498,-.7497] |
| AV45*HippoV | 0 | -.1692 | -.0001 |
|  | [0,0] | [-.1694,-.1691] | [-.0002,0] |
| CSF tau*HippoV | 0 | -.0807 | -.0004 |
|  | [0,0] | [-.0808,-.0806] | [-.0005,-.0003] |

### Agreement between empirical and simulated data

The estimated standardized coefficients of simple regression analyses between real and simulated data were .823, .960, .934 and .757 for amyloid PET, CSF tau, hippocampal volume and ADAS-cog, respectively. The corresponding adjusted *R*^2^ values were .68, .90, .84 and .53. The simulated values thus showed substantial agreement with real data. See figure 1 for plots of simple regression analyses.

**Figure 1.**
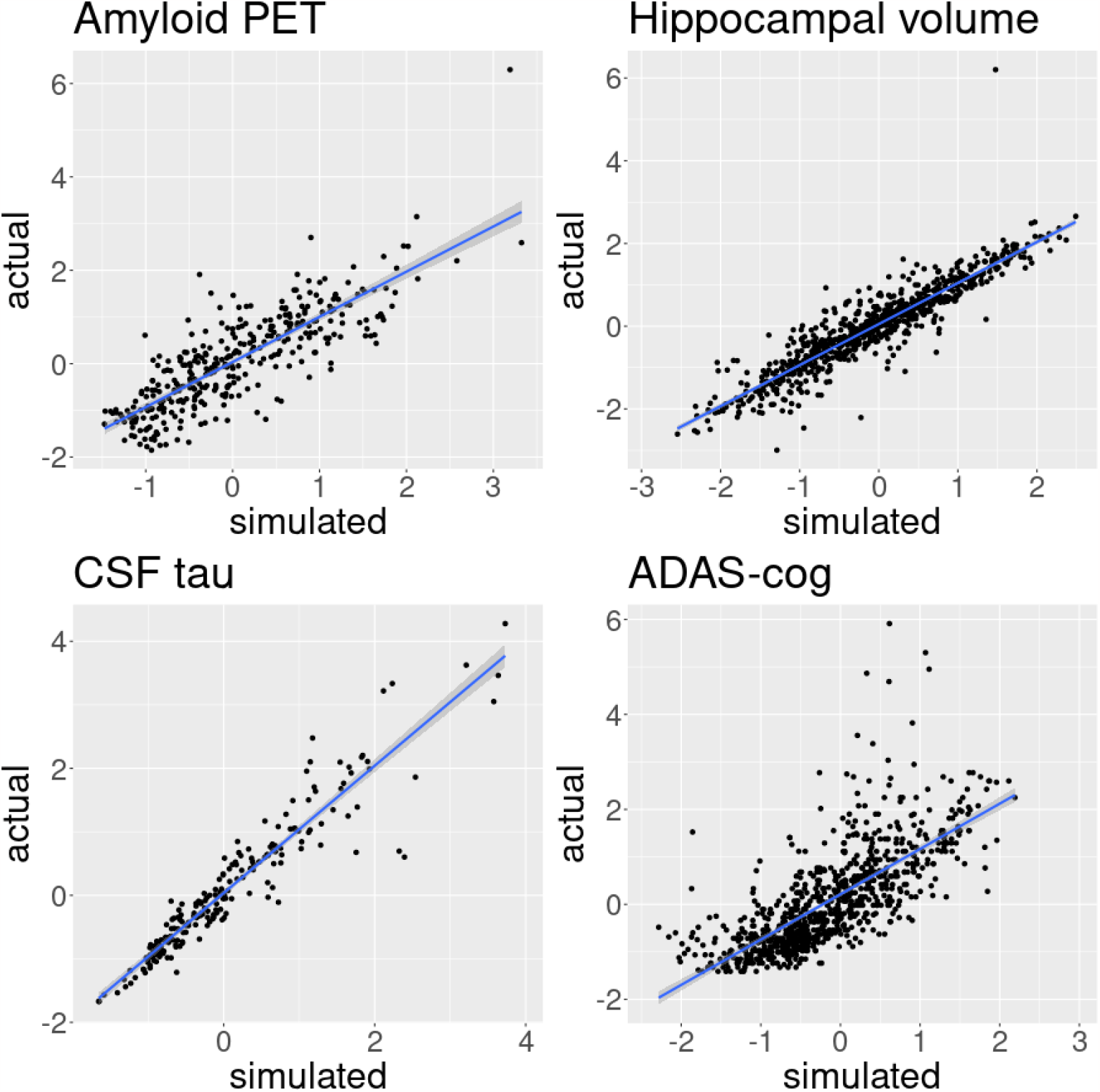
Actual versus simulated biomarker and cognitive outcome data points. This plot shows the empirical data for all subjects at all available timepoints versus the corresponding simulated data points as predicted from the mathematical model with a line representing simple linear regression. CSF tau, corticospinal fluid total tau. ADAS-cog, cognitive Alzheimer’s Diseas Assessment Scale.

### Dependence of interaction effects on data sampling

Regression models yielded estimates of comparable magnitude for main effects (i.e. biomarkers) on cognitive outcome across the different simulated data samples. Specifically, for cross sectional sampling that was consistent with respect to the time point of the data, the average standardized regression coefficients were .3369, .1679 and -.8710 for amyloid-PET, CSF tau and hippocampal volume, respectively. The corresponding 95% confidence intervals were [.3171,.3567], [.1665,.1692] and [-.8731,-.8690]. For cross sectional sampling that was random with respect to the time point of the data, average standardized regression coefficients were .2627, .1286 and -.7586 for amyloid-PET, CSF tau and hippocampal volume, respectively. The corresponding 95% confidence intervals were [.2626,.2627], [.1286,.1287] and [-.7587,-.7585]. For cross sectional sampling that was random with respect to the time point of the data and with the effect of time regressed out of main effects, average standardized regression coefficients were .2987, .1448 and -.7498 for amyloid-PET, CSF tau and hippocampal volume, respectively. The corresponding 95% confidence intervals were [.2986,.2988], [.1447,.1449] and [-.7498,-.7497].

However, interaction effect estimates differed between simulated data samples. Specifically, for cross sectional sampling that was consistent with respect to the time point of the data, average standardized regression coefficients were 0 for interactions of amyloid-PET and CSF tau with hippocampal volume. In contrast, for cross sectional sampling that was random with respect to the time point of the data, average standardized regression coefficients were -.1692 and -.0807 for amyloid-PET with hippocampal volume and CSF tau with hippocampal volume, respectively. The corresponding 95% confidence intervals were [-.1694,-.1691] and [-.0808,-.0806]. These interaction effects were such that at higher levels of hippocampal volume the detrimental effect of high amyloid-PET and CSF tau levels on cognitive outcome was attenuated. However, these interaction effects decreased by several orders of magnitude when repeating the random sampling but regressing the effect of time out of main effects prior to estimating regression models. The average standardized regression coefficients for this analysis were -.0001 and -.0004 for amyloid-PET with hippocampal volume and CSF tau with hippocampal volume, respectively. The corresponding 95% confidence intervals were [-.0002,0] and [-.0005,-.0003]. See Table 3 for an overview and figure 2 for boxplots of estimated regression coefficients.

**Figure 2.**
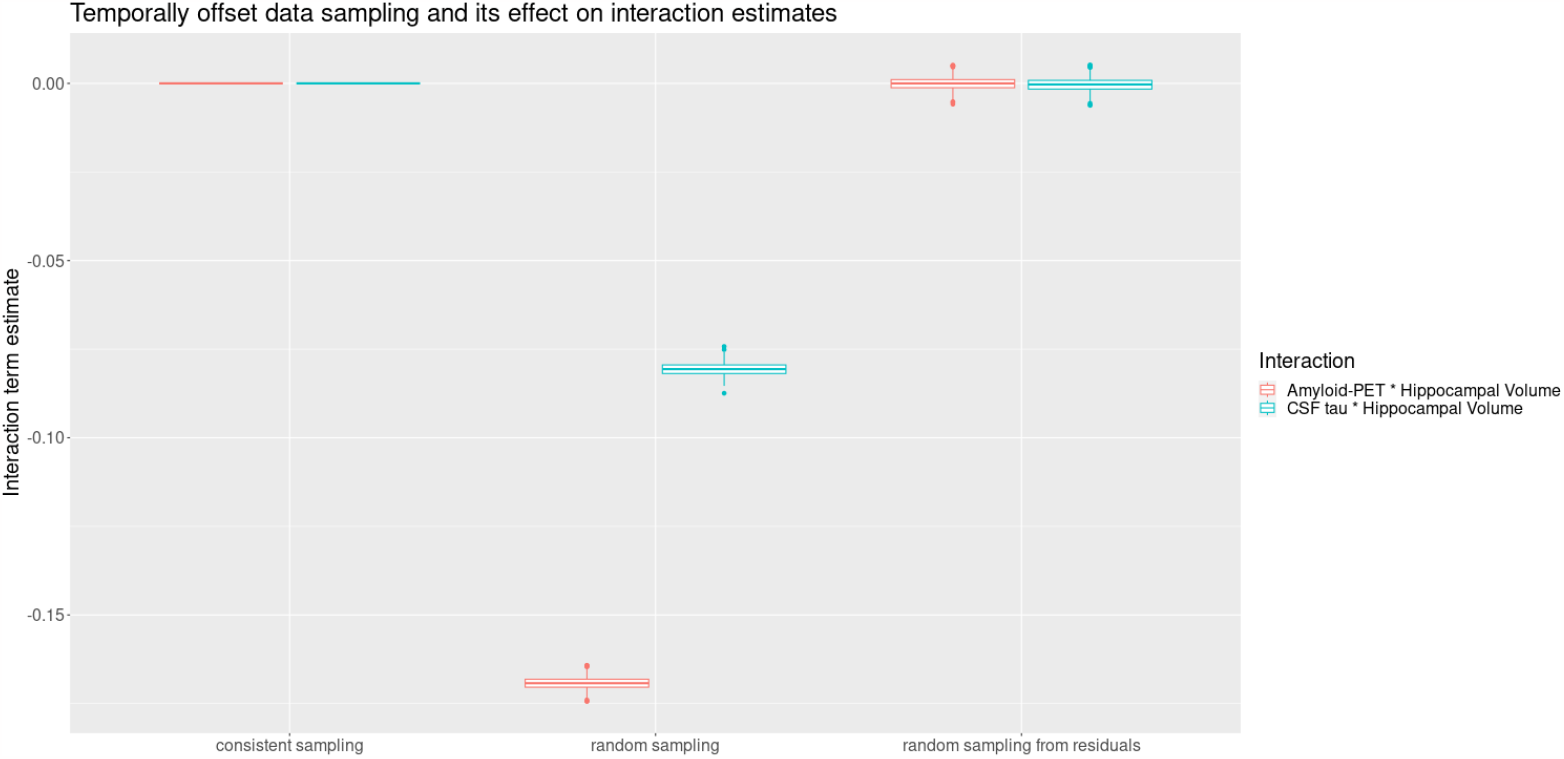
Effect of temporally offset data sampling on interaction term estimates. Shown are boxplots of interaction term estimates from regression analyses on simulated data, where individual trajectories of biomarkers and cognition differed only due to initially varied states of interaction term main effect variables. Consistent sampling, data points were sampled cross sectionally from one corresponding timepoint after baseline from each individual trajectories. Random sampling, data points were sampled from a random timepoint cross sectionally from each individual trajectory. Random sampling from residuals, as in random sampling but with the variance associated with time after baseline removed from main effect variables.

## Discussion

The results of the present study demonstrate how interaction effects in regression analyses can arise as a statistical artifact due to cross sectional sampling. The faux interaction effects were demonstrated using simulated data that was generated by a mathematical AD biomarker cascade model, whose parameters were estimated from longitudinal empirical data and whose predictions showed substantial agreement with empirical data points. Furthermore, the faux interaction effects were greatly reduced when regressing out the variance associated with time point of sampling from main effect variables. This finding has important implications for the inferences drawn from interaction effects that could be deemed to indicate cognitive reserve. To our knownledge, this issue has not been extensively discussed prior to this publication.

### Model assessment

Previous studies have employed similar methods, i.e. differential equation models, to model AD. However, some of these studies focused on specific aspects of the disease process whereas most arguably used conceptually as well as mathematically more elaborate models [4, 12, 19–21]. In contrast, for the present study, we aimed to demonstrate a statistical artifact through a mathematical model that was as simple as possible in order to be able to argue that it may be present in more complex conceptions of AD as well.

The mathematical model’s parameters, i.e. the associations between biomarkers and cognitive reserve as well as their change over time, were estimated from empirical data and are generally in agreement with prior studies. Specifically, the deterioration of all biomarkers over time for patients of AD, i.e. increase of cerebral amyloid and CSF tau, hippocampal atrophy and cognitive decline, has been demonstrated repeatedly.

The agreement of the simulated data points produced by the mathematical model and corresponding empirically available data points was substantial both for biomarkers as well as cognitive outcome, as indicated by standardized regression coefficients ranging between .746 and .967. As the corresponding *R*^2^ values ranged between .54 and .90, we believe the model captures the majority of the temporal evolution of the biomarkers and cognitive outcome considered within the sample and the available time span from 6 to 186 months after baseline.

### Faux interaction effects

The mathematical model was explicitly designed not to contain a modulation of biomarkers’ associations with cognitive outcome by other biomarkers. Instead, the mathematical model of the AD biomarker cascade consisted entirely of independent associations between biomarkers and the rate of change of other biomarkers or cognitive outcome. These associations were estimated from empirical data (refer to equation (2) and Table 2). Correspondingly, regression analyses on simulated data, where biomarker levels and cognitive outcome had been estimated at a specific time point after baseline from initial biomarker levels individually and randomly varied across their empirical ranges, showed significant independent main effects of biomarkers on cognitive outcome but no interaction between biomarkers (see Table 3).

However, in naturalistic cross sectional designs, individually varying levels of biomarkers at the beginning of the disease process are confounded with a measurement time point relative to the beginning of the disease process, which is therefore difficult or impossible to estimate and take into account. This circumstance was integrated into the above analysis by randomly varying the timepoint after baseline at which the mathematical model simulated biomarker levels and cognitive outcomes. The consequent appearance of interaction effects between biomarkers on cognitive outcome when repeating regression analyses demonstrates that when the measurement time point relative to the disease process is not taken into account, interaction effects between biomarkers may arise purely as a statistical artifact. Indeed, they are greatly diminished when regressing the effect of time after baseline out of variables and repeating this analysis.

The mechanism by which these faux interaction effects arise is the following. For the simulated data produced by the mathematical model, the following can be said to be generally true based on the estimated parameters shown in Table 2. First, cognitive outcome will be lower at later points in time after baseline. Second, all biomarkers will also be at worse levels at later time points after baseline. Third, this implies that data points where one biomarker is more altered than the others are at earlier time points relative to baseline. A design where one measures biomarkers and cognitive outcome at a random time point for each individual’s trajectory would consequently yield the following statistical observations for an interaction analysis disregarding the time points of measurements: data points where only one of the main effect variables are altered correspond to better cognitive performance, because they are earlier in time. Data points, where both main effects are altered would correspond to worse cognitive performance than explained by the biomarkers’ individual effects due to being later in time relative to baseline. In other words, time after the beginning of the disease process confounds the interaction term of biomarkers. Please refer to figures 3 and 4 for a visualization.

**Figure 3.**
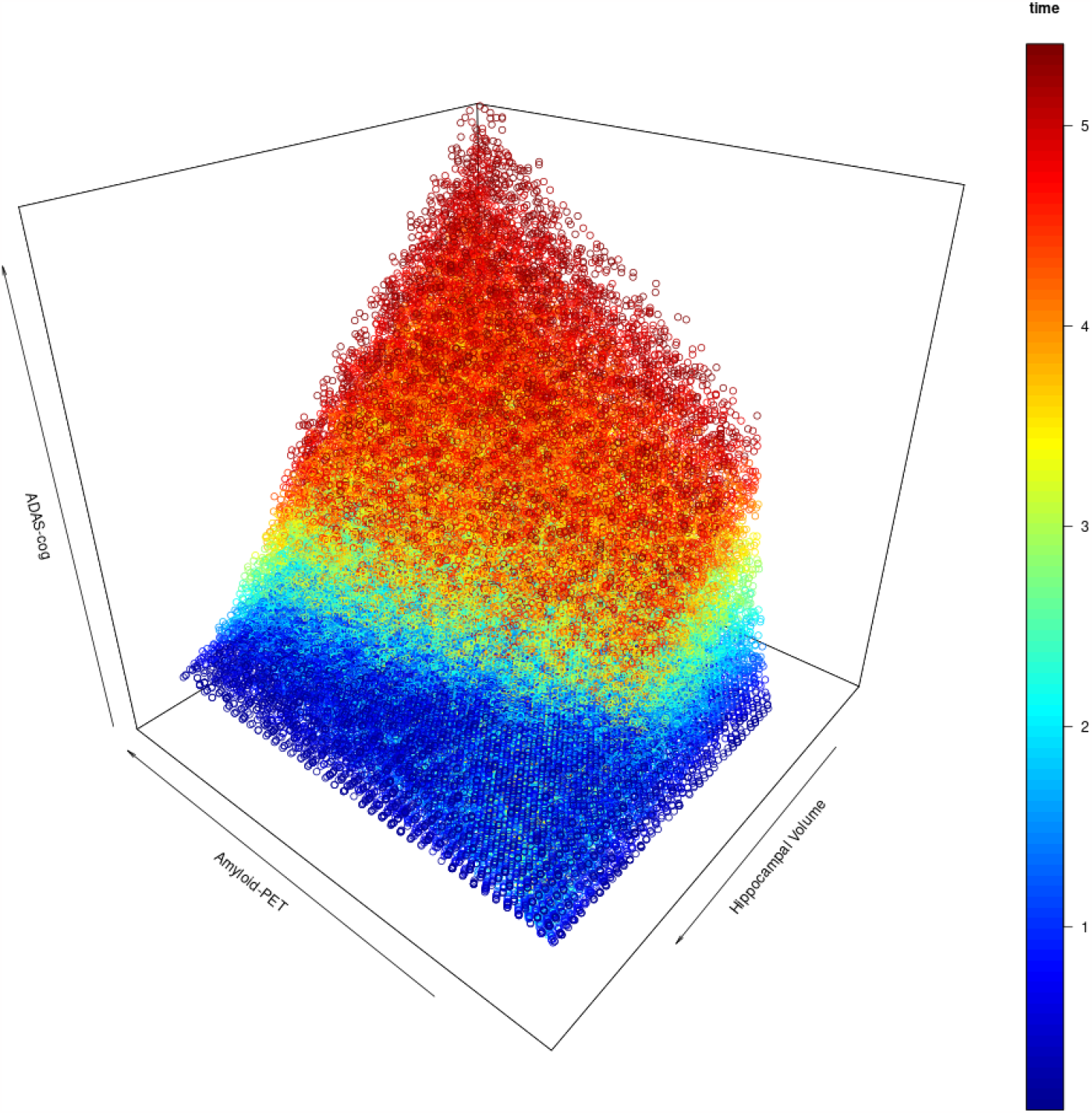
Simulated data points for amyloid-PET and hippocampal volume. Scatter plot of simulated data points for initial configurations of amyloid-PET and hippocampal volume varying between -2 and 2 standard deviations from the mean of their empirical distribution. Simulation time after baseline in years is color coded. At baseline, all simulated subjects start at the empirical mean cognitive outcome, amyloid and hippocampal volume are distributed independently. At later time points, cognitive outcome tends to be worse and amyloid-PET and hippocampal volume are jointly altered under the AD biomarker cascade model. Randomly sampling one time point for each simulated subject from this distribtion may lead to the estimation of interaction effects between amyloid-PET and hippocampal volume that are driven by the underlying temporal dynamic rather than cognitive reserve. ADAS-cog, Alzheimer’s Disease Cognitive Assessment Scale.

**Figure 4.**
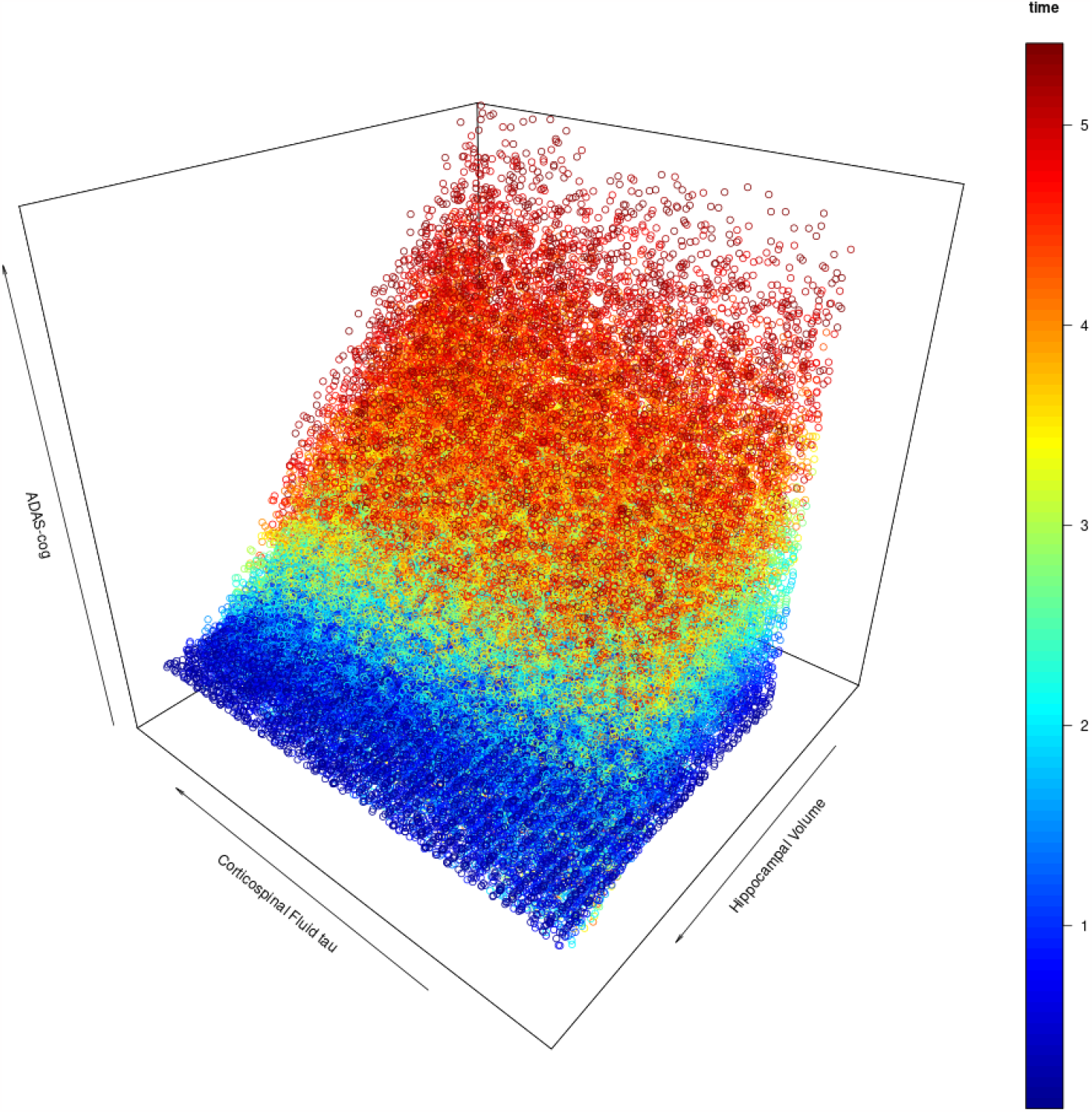
Simulated data points for corticospinalfluid tau and hippocampal volume. Scatter plot of simulated data points for initial configurations of corticospinal fluid tau and hippocampal volume varying between -2 and 2 standard deviations from the mean of their empirical distribution. Simulation time after baseline in years is color coded. At baseline, all simulated subjects start at the empirical mean cognitive outcome, tau and hippocampal volume are distributed independently. At later time points, cognitive outcome tends to be worse and tau and hippocampal volume are jointly altered under the AD biomarker cascade model. Randomly sampling one time point for each simulated subject from this distribtion may lead to the estimation of interaction effects between tau and hippocampal volume that are driven by the underlying temporal dynamic rather than cognitive reserve. ADAS-cog, Alzheimer’s Disease Cognitive Assessment Scale.

### Conclusions for the inference on cognitive reserve

Statistical interaction effects have been proposed and used to identify biomarkers of neurophysiological cognitive reserve surrogates, that facilitate functional adaption to the presence of pathology [29]. However, under the premise of the AD biomarker cascade model, statistical interaction effects between biomarkers may arise as an artifact of the data sampling strategy. Specifically, the levels of biomarkers, both of pathology or candidate surrogates of cognitive reserve, and cognitive outcome are a convolute of individually differing initial predispositions (potentially representing brain maintenance and brain reserve) and the amount of time that they have been affecting each other according to the AD biomarker cascade model. Even though associations among biomarkers and cognitive outcome are constant under the biomarker cascade model, data points from later in the disease process will show stronger conjoint alterations of biomarkers and cognitive outcome and may thus drive statisticall estimated interaction effects when mixed with data points from earlier in the disease process in cross sectional designs. For individual data points, time spent under the regime of the biomarker cascade model’s associations among biomarkers and cognitive outcome thus should to be incorporated in some form into statistical analyses.

### Limitations

This study has several limitations, mainly regarding the mathematical model, which could be much more enhanced with respect to biomarkers or the associations among them. Additionally, the sample from which the data has been estimated may not accurately represent the population due to its recruitment schemes. The initial configurations of the simulated data do not correspond to an empirical joint distribution of the respective biomarkers. However, this study’s aim is not to make any inferences with regard to the actual disease process of AD nor the prediction of biomarker statuses. Rather, the aim was to demonstrate in a proof-of-concept fashion with a minimal AD mathematical model that misleading statistical interaction effects can arise under the very premise of the core of the AD cascade model.

## Acknowledgments

We thank Tatjana Tchumatchenko of the Institute of Physiological Chemistry at the University Medical Center Mainz for her aid and insights during the preparation of the manuscript.

Data collection and sharing for this project was funded by the Alzheimer’s Disease Neuroimaging Initiative (ADNI) (National Institutes of Health Grant U01 AG024904) and DOD ADNI (Department of Defense award number W81XWH-12-2-0012). ADNI is funded by the National Institute on Aging, the National Institute of Biomedical Imaging and Bioengineering, and through generous contributions from the following: AbbVie, Alzheimer’s Association; Alzheimer’s Drug Discovery Foundation; Araclon Biotech; BioClinica, Inc.; Biogen; Bristol-Myers Squibb Company; CereSpir, Inc.; Cogstate; Eisai Inc.; Elan Pharmaceuticals, Inc.; Eli Lilly and Company; EuroImmun; F. Hoffmann-La Roche Ltd and its affiliated company Genentech, Inc.; Fujirebio; GE Healthcare; IXICO Ltd.; Janssen Alzheimer Immunotherapy Research & Development, LLC.; Johnson & Johnson Pharmaceutical Research & Development LLC.; Lumosity; Lundbeck; Merck & Co., Inc.; Meso Scale Diagnostics, LLC.; NeuroRx Research; Neurotrack Technologies; Novartis Pharmaceuticals Corporation; Pfizer Inc.; Piramal Imaging; Servier; Takeda Pharmaceutical Company; and Transition Therapeutics. The Canadian Institutes of Health Research is providing funds to support ADNI clinical sites in Canada. Private sector contributions are facilitated by the Foundation for the National Institutes of Health (www.fnih.org). The grantee organization is the Northern California Institute for Research and Education, and the study is coordinated by the Alzheimer’s Therapeutic Research Institute at the University of Southern California. ADNI data are disseminated by the Laboratory for Neuro Imaging at the University of Southern California.

